# ISGylation drives basal breast tumour progression by promoting EGFR recycling and Akt signalling

**DOI:** 10.1101/767533

**Authors:** Alfonso Bolado-Carrancio, Morwenna Muir, Ailith Ewing, Kenneth Macleod, William Gallagher, Lan Nguyen, Neil Carragher, Colin Semple, Valerie G Brunton, Patrick Caswell, Alex von Kriegsheim

## Abstract

ISG15 is an ubiquitin-like modifier that is associated with reduced survival rates in breast cancer patients. However, the mechanism by which ISG15 achieves this remains elusive. We demonstrate that modification of Rab GDP-Dissociation Inhibitor Beta (GDI2) by ISG15 (ISGylation) alters endocytic recycling of the EGF receptor (EGFR). By regulating EGFR trafficking, ISGylation sustains Akt-signalling *in vitro* and *in vivo*. Persistent and enhanced Akt activation explains the more aggressive tumour behaviour observed in animal models and human breast cancers. We show that ISGylation can act as driver of tumour progression rather than merely being a marker of it.

## Introduction

Interferon-Induced 15 kDa protein (ISG15) was the first ubiquitin-like protein identified, but its research was long restricted to the field of immune response where it was initially discovered. Recently, ISG15 has been associated with processes and pathologies distinct from the innate-immune response (Villarroya-Beltri, Guerra et al., 2017). Tumour progression and aggressiveness of several cancer types, including endometrium, bladder, prostate, melanoma, colorectal, liver and breast cancer (Andersen, Aaboe et al., 2006, Kiessling, Hogrefe et al., 2009, Li, Wang et al., 2014, Padovan, Terracciano et al., 2002, Talvinen, Tuikkala et al., 2006); (Bektas, Noetzel et al., 2008, Nabet, Qiu et al., 2017, Weichselbaum, Ishwaran et al., 2008) has been correlated to ISG15 expression. Especially in breast cancer, ISG15 is considered a potential biomarker. However, the mechanism through which ISG15 regulates tumorigenesis remains elusive.

ISG15, like other ubiquitin-like proteins, can be covalently bound to lysine residues of target proteins (Loeb & Haas, 1992) in a process known as ISGylation. This post-translational modification (PTM) is similar to ubiquitination, as it requires the sequential co-ordination of a cascade involving three different ligases. The process can be reversed by the action of Ubiquitin Specific Peptidase 18 (USP18), a member of the deubiquitinase family which is the only ISG15-specific deubiquitinase enzyme that has been described so far (Malakhov, Malakhova et al., 2002).

Basal levels of ISG15 and ISGylation-related enzymes are generally low in cells. Protein expression can be substantially increased by a variety of stimuli, such as type I interferons (Korant, Blomstrom et al., 1984), Lipopolysaccharide (LPS) (Malakhova, Malakhov et al., 2002), growth factors (Farrell, Kelly et al., 2014) or viral infections (Haas, Ahrens et al., 1987). In breast cancer cells, enhanced ISG15 expression is induced through exosome-mediated cGAS activation (Boelens, Wu et al., 2014, Nabet et al., 2017) and from nuclear DNA release after DNA damage (Mackenzie, Carroll et al., 2017).

ISG15 functions either as unconjugated or as a covalently linked protein. When released into the extracellular matrix (Knight & Cordova, 1991), it functions as an immunomodulatory agent for different cell types, such as lymphocytes (Villarreal, Wise et al., 2015) or Natural Killer (NK) cells (D’Cunha, Ramanujam et al., 1996). The role of secreted ISG15 during tumour development is contradictory. It has been described both as an antitumoural factor by increasing NK cell infiltration in xenografts (Burks, Reed et al., 2015) or tumourigenic by increasing the invasive potential of primary tumour cells (Sainz, Martin et al., 2014). As a conjugate, ISG15 is also widely linked to immune functions.

Initially it was primarily considered an antiviral protein (Morales & Lenschow, 2013), although recent data in human models suggest that ISG15 can function as a negative effector of type I interferon signalling rather than directly regulating functions of the immune system (Hermann & Bogunovic, 2017).

To comprehensively map the role of ISGylation in both the innate immune response and tumorigenesis, numerous mass spectrometry-based studies have been carried out to identify substrates of ISG15 modification, the ISGylome, as defined by the proteins which are ISGylated in a particular system (Giannakopoulos, Luo et al., 2005, Malakhov, Kim et al., 2003, Zhao, Denison et al., 2005). These studies showed that putative ISG15 substrates are associated with multiple cellular signalling pathways and are cell/tissue type dependent, thus suggesting that ISG15 might play a broader role in the cell and not just during the interferon-associated response. However, only a few of these putative targets have been validated endogenously and even fewer have been functionally characterised by identifying the molecular role of the ISGylation. Therefore, our current knowledge of how ISGylation affects cellular function and signalling is limited. This is a challenge that needs to be overcome before we can elucidate the role of ISG15 in tumorigenesis. Despite the limited number of characterised substrates, it is apparent that the molecular function of ISGylation is varied. It has been shown to induce both protein stabilization (Desai, Haas et al., 2006) and degradation (Burks, Reed et al., 2014, Huang, Wee et al., 2014), as well as modulating protein-protein interactions (Cerikan, Shaheen et al., 2016, Jeon, Choi et al., 2009). This plasticity, reminiscent of ubiquitination, makes ISG15 a dynamic, reversible post-translation modification that can apparently regulate substrates disparately.

In breast cancer models, ISG15 and/or ISGylation correlate with aggressive tumorigenic features such as, cell cycle progression, cell motility and tumour growth in xenograft models (Burks et al., 2014, Desai et al., 2006, Desai, Reed et al., 2012). In these models, ISGylation has been suggested to behave as a competitive inhibitor of ubiquitination by binding and masking potential ubiquitination sites. However, little direct mechanistic insight on how ISG15/ISGylation elicit these functions has been gained.

ISG15 has been broadly associated with breast cancer progression (Chaohui Zuo 2016), nevertheless, we still do not know if ISG15 is just a biomarker or indeed a driver of aggressive tumour and metastatic behaviour. We and others have discovered that ISG15 can induce the migratory potential by supporting epithelial-mesenchymal transition (Burks et al., 2014, Farrell et al., 2014) and there is strong correlation between ISG15 expression and breast cancer progression. However, significant progress in uncovering a functional link between cause and effect has been hampered by the lack of functional, mechanistic insight into how ISGylation regulates signalling networks. We decided to bridge this gap by applying unbiased, systems approaches to elucidate in detail how ISGylation contributes to cell signalling.

In this study, we aimed to identify the molecular mechanisms that explains why high ISG15/ISGylation correlates with poor patient prognosis in breast cancer.

## Results

### ISGylation negatively correlates with disease-free survival

We first used the Breastmark mRNA database (Madden, Clarke et al., 2013) to analyse the correlation between prognosis, ISG15 levels and metastasis. As previously shown, elevated ISG15 mRNA levels correlated with lower disease-free survival (Fig.1A). Surprisingly however, this correlation was lost in the patient cohort where the tumour had not spread to the lymph nodes (Fig.1B). The correlation was only maintained in patients with identified lymph node metastasis (Fig.1C). After separating the patients into breast cancer subgroups (Luminal A/B, HER2+, and Basal), analysis of disease-free survival suggested a modest correlation between ISG15 and Luminal B and Basal subtypes (Fig. 1D). This analysis also suggests that overexpression of HER2 and ISG15 are not additive and that both induce overlapping signalling pathways beneficial for tumour progression required in either of the other non-HER2-overexpressing subgroups.

**Fig 1.**
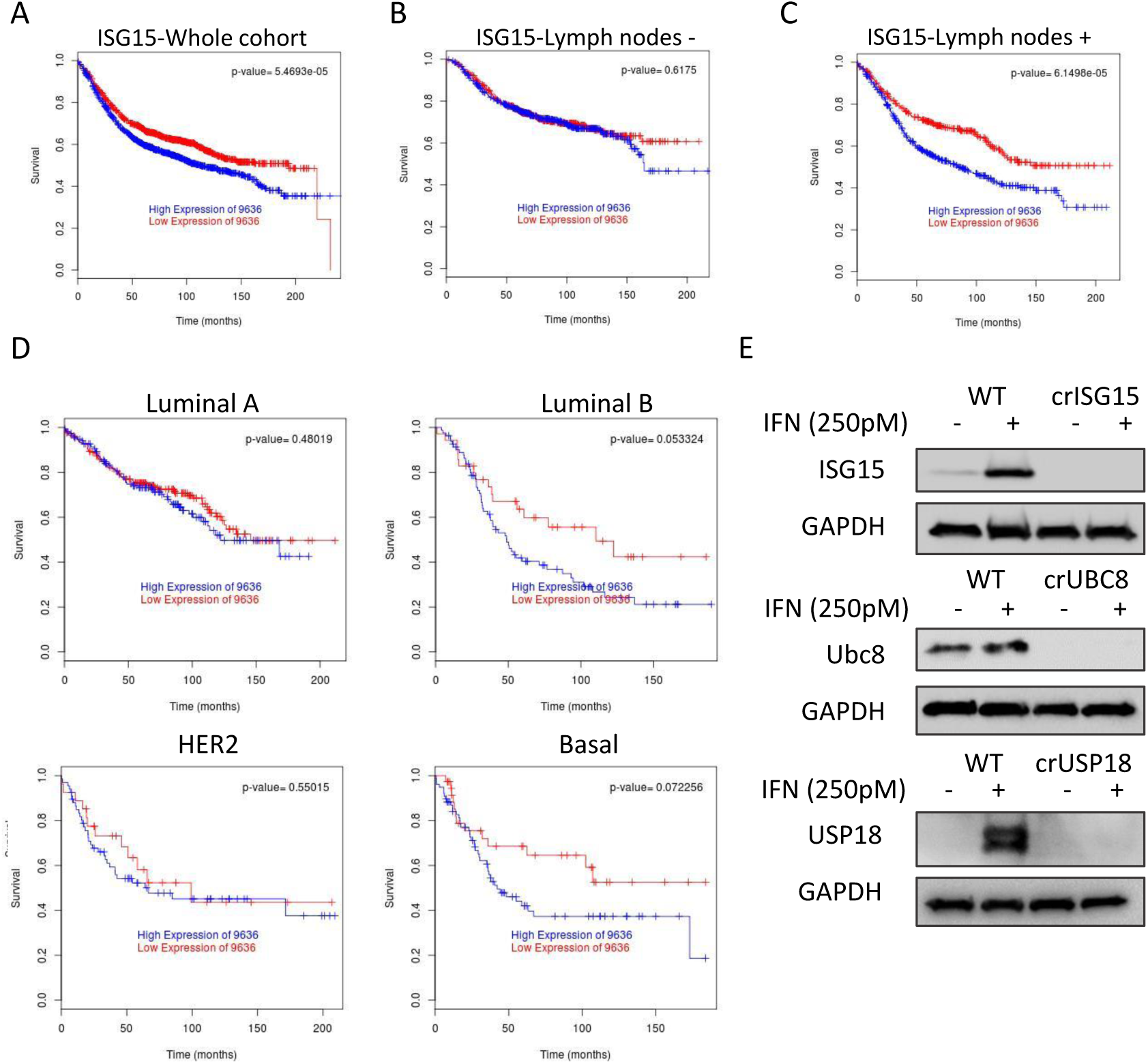
ISG15 expression is associated with poor outcome in breast cancer patients with lymph-node metastasis. (A) Kaplan–Meier plot of disease-free survival of breast cancer patients (n=2643) separated into two groups with high (blue) or low (red) mRNA expression levels of ISG15. (B) Kaplan-Meier plot of free disease survival of patients from A that are negative for tumour cells in lymph nodes (n=740) (C) Kaplan-Meier plot of free disease survival of patients from A positive for tumour cells presence in lymph nodes (n=1178). (D) Kaplan-Meier plot of cohort displayed in (A) classified by breast cancer subtype: luminal A (n= 366), luminal B (n=117), HER2 (n=94), basal (n=114). (E) WB analysis of the indicated representative clones to confirm the knockout, cells were treated for 48h with either vehicle or IFNb1a 250pM to boost the protein levels of the different ISGylation-related proteins.

ISG15 can function as a conjugated or free/secreted protein. Therefore, to narrow down which form is associated with this correlation, we wanted to determine if regulators of ISGylation showed an analogous correlation. Thus, we also analysed the correlation between UBE2L6, which is the gene that encodes the main ISG15 E2 ligase UBC8, and disease-free survival. As seen with ISG15, UBE2L6 mRNA expression levels were negatively correlated with survival in patients with cancer that had spread to the lymph nodes (Fig.S1A) but the correlation was lost in patients without lymph node metastasis (Fig.S1B). When separating the cohort by cancer subtypes, a statistically significant correlation was found in the basal subtype (Fig.S1C).Together, these data strongly suggest that UBC8-mediated ISG15 conjugation is elemental for the identified association between ISG15 and disease-free survival in lymph-node positive patients.

### ISGylation enhances cellular aggressiveness

The inverse correlation between ISG15 or UBC8 expression in lymph node metastasis positive patients with disease-free survival suggested that the conjugated ISG15 rather than the free/secreted protein enhances metastasis and tumour progression. To study the likely association between ISGylation and breast cancer metastasis, we generated different cell line models with varying levels of ISGylation in MDA-MB-231-luc-D3H2LN. Using CRISPR/Cas9 and two specific gRNAs per gene we either knocked-out ISG15 (crISG15), UBE2L6 (crUBC8), USP18 (crUSP18) or generated a control line with the same Cas9 expression plasmid but without targeting gRNA (WT). These four cell lines allowed us to test the molecular and cellular characteristics of cells devoid of ISG15 (crISG15), devoid of ISGylation (crUBC8), with enhanced levels of ISGylation (crUSP18) or with basal levels of ISG15 and ISGylation (WT). Confirmation of the respective knockouts was performed by western blot (Fig.1E). Due to the low basal levels of ISG15, USP18, UBC8 and ISGylation we additionally incubated the cells with IFN1b 250pM (or vehicle) to enhance expression levels and facilitate the confirmation of protein depletion. Changes of ISGylation were confirmed by western blot, in the presence of either vehicle or IFN1b for 48h (Fig.2A). The result confirmed that crISG15 expressed no free or conjugated ISG15, crUBC8 expressed only unconjugated ISG15 and clones of crUSP18 enhanced the levels of ISGylation to such an extent that numerous protein band appeared to be prominently modified even under basal conditions.

**Fig 2.**
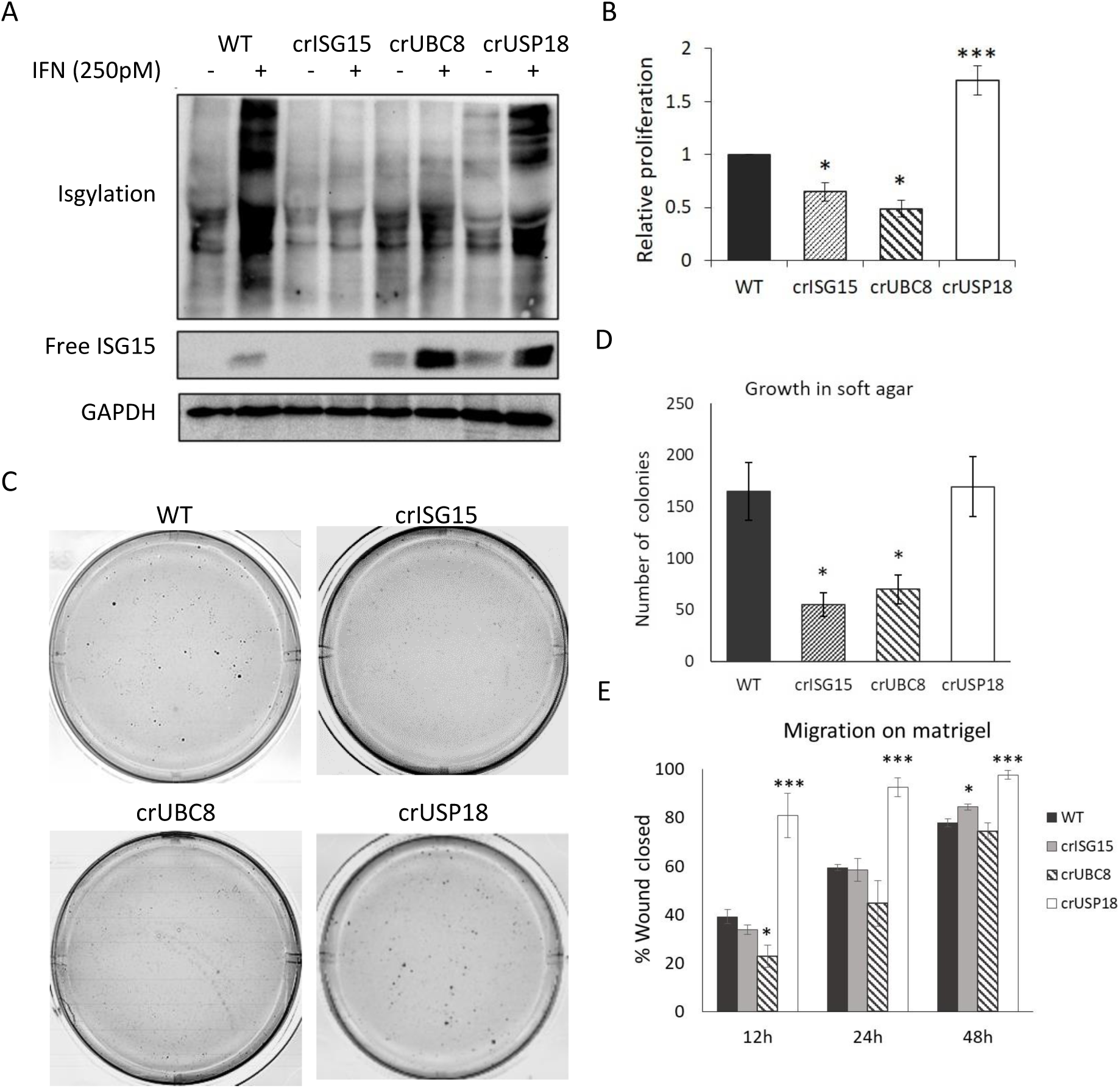
ISGylation correlates with an aggressive phenotype in MDA-MB-231-luc-D3H2LN. (A) WB analysis of free ISG15 and ISGylation levels in WT cells and the different representative clones, treated for 48h with either vehicle or IFNb1a 250pM. (B) Proliferation assay performed in the indicated clones. Equal amounts of cells were seeded and then counted manually. Bar graph shows the relative increase in cell number during 72h ± S.E.M. n=4. (C) Representative images of soft agar experiments. Equal cell numbers of the indicated, representative clones were seeded embedded in soft agar and after 28 days stained with crystal violet; n=4. Images are displayed in grayscale for better identification of the colonies. (D) Quantification of number of colonies identified in the soft agar assay. Bar graph shows average ± S.E.M. (E) Relative migration. Equal amounts of cells were seeded into matrigel coated wells where scratch wounds was used to determine cell motility, photos were taken every 3h for up to 48h and motility was measured as percentage of wound closed per time point. Bar graph shows the average migration of the different clones versus WT cells, at 12, 24 and 48h, ±S.E.M; n=3. p-value < 0.05 (*), p-value < 0.005 (***).

Having generated these models, we assessed if cellular phenotypes associated with tumour aggressiveness could be linked to ISGylation. We were particularly interested in cellular attributes linked to tumour progression. Consequently, we focused on proliferation, anchorage-independent growth and cell motility or invasion into extracellular-like matrix. We found a positive correlation between ISGylation and the proliferation rate (Fig.2B), the ability to form colonies in soft agar (Fig.2C); the cell-density of those colonies (Fig.S2A) and their total number (Fig.2D). Using the different clones and in the presence or absence of EGF or serum, we assessed the migration potential of the individual cell lines in a wound-healing assay. In this assay only crUSP18 cells, showed a statistically significant increase in motility (Fig.2E). When assaying invasion into matrigel, crUSP18 statistically increased the ability of cells to invade. In addition, we observed a trend indicating that cells devoid of ISGylation, crISG15 and crUBC8, had a reduced ability to invade (Fig.S2B and S2C). Overall, these data suggests that *in vitro* ISGylation increases several markers of tumour aggressiveness in a basal breast cancer cell line.

### ISGylation enables sustained Akt-signalling

Depending on the context, ISG15 has been reported to regulate multiple signalling pathways, including Akt (Yanguez, Garcia-Culebras et al., 2013), ERK (Malakhov et al., 2003) or JAK/STAT (Malakhova, Yan et al., 2003). Therefore, to determine which, if any, signalling networks are regulated by ISG15/ISGylation in MDA-MB-231 cells, we employed an unbiased, systematic approach to quantify pathway activities. Using a Reverse Phase Protein Array (RPPA) we monitored how expression and phosphorylation of 58 signalling proteins were regulated at basal levels and upon EGF stimulation in the CRIPSR cell line panel. RPPA covered major kinase signalling pathways and allowed us to determine which networks were affected by varying levels of ISGylation. We stimulated the panel of CRISPR clones for 0, 10, 30 or 60 min with EGF (Fig.3A). Despite not detecting changes in EGFR levels or phosphorylation across the clones, we identified pathways that were regulated by ISGylation. Cells lacking ISG15 activated STAT1, an observation in agreement with previous reports (Sooryanarain, Rogers et al., 2017). In addition, we observed the increased expression of MAPKAPK2, a kinase downstream of p38 (Krump, Sanghera et al., 1997). Neither STAT1 nor MAPKAPK2 were altered in crUBC8 cells, it is therefore unlikely that ISGylation regulates these pathways. We further detected a subtle reduction in ppERK levels in both crISG15 and crUBC8 at 10min when compared to WT and crUSP18. The pathway most strikingly affected by ISGylation among all conditions was PI3K/Akt. pAktser473 (pAkt) levels were positively correlated with ISGylation levels, with a maximal reduction of pAkt levels at 10 minutes EGF stimulation in the crISG15 and crUBC8 compared to the control (Fig.3B). Additionally, we observed higher peak and sustained Akt phosphorylation in crUSP18 clones at 10 and 30min. We confirmed the observed regulation of Akt and downstream pathway by western blot (Fig.3C). We further validated the association between pAkt and ISGylation by transiently transfecting an ISG15 expressing plasmid into the cell lines. After stimulation with EGF for 10min. in crISG15 and WT cells re- or overexpression of ISG15, respectively, rescued or increased pAkt levels, whereas ISG15 overexpression did not alter pAkt in crUBC8 (Fig.S3A). These rescue experiments demonstrated that sustained Akt signalling is enhanced by ISGylation rather than unconjugated ISG15. Neither RPPA nor the western blot data (Fig. 3A and B) indicated that decreased Akt activation was due to altered EGFR expression or phosphorylation.

**Fig. 3.**
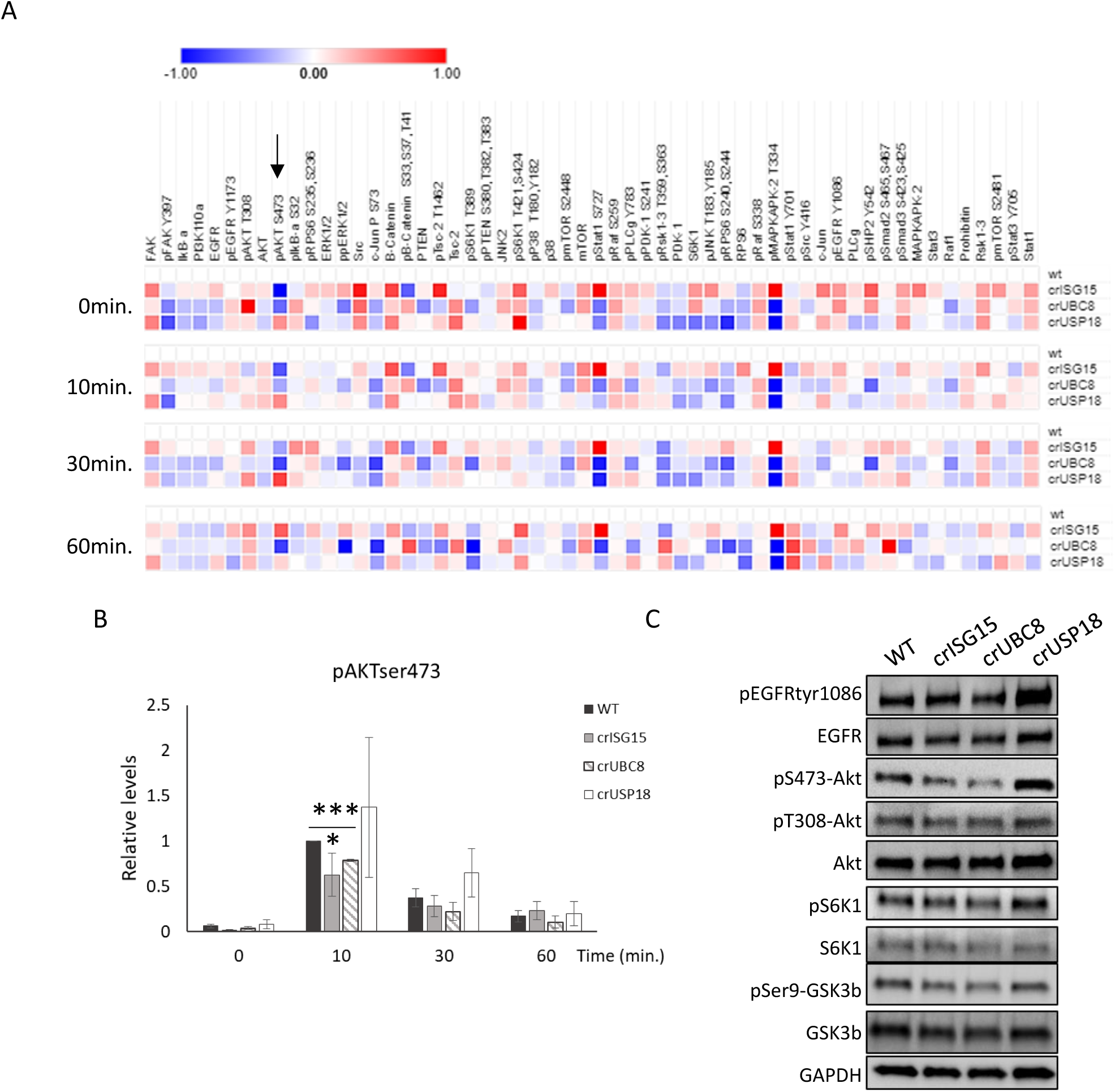
Loss of ISGylation is associated with decreased Akt phosphorylation. (A) RPPA analysis performed in WT and the different ISGylation CRISPR clones, treated with EGF 10ng/ml for 0, 10, 30 and 60 minutes. Heatmap displays the Log2 of intensity average of biological triplicates normalized to the WT one at the different time point. Arrow indicates pAktser473 data. n=3 (B) Bar graph shows the values for pAktser473 obtained in the RPPA. Values displayed are the mean ±S.E.M. (C) Confirmation of RPPA data by WB analysis of Akt pathway activation in the different clones treated with EGF 10ng/ml for 10 minutes. p-value < 0.05 (*).

The lack of an ISG15-dependent effect on EGFR signalling upstream of Akt, with no differences observed in EGFR, pEGFR, Akt or PI3K protein levels (Fig.3A), suggested that ISGylation might regulate feedback loops that would alter pAkt dynamics. To confirm that the observed suppression of Akt phosphorylation upon ISGylation-loss was independent of EGFR, WT and crISG15 cells were treated with several concentrations of insulin, a strong activator the PI3K/Akt pathway, which signals through the insulin receptor. Similar to our observation with EGF, pAktser levels were suppressed in crISG15 compared to WT (Fig.S3B).

To determine if Akt suppression was due to altered feedback/network regulations, we performed extended time course experiments in the different clones, stimulating the cells with EGF for 0, 2, 5, 10, 30 and 60 minutes. Our results (Fig.S3C and D) indicated that at 5 minutes peak pAkt levels were similar. However, after the early time points, pAkt levels diverged depending on the ISGylation status, with pAkt suppressed at 10, 30 and 60 minutes post-stimulation in clones lacking ISGylation. This suggests that the signalling sustenance rather than upstream and immediate activation, is influenced by ISGylation.

### ISGylation is required for efficient receptor recycling

ISGylation has been shown to regulate PTEN stability (Mustachio, Kawakami et al., 2017). In our system we were unable to detect any regulation of PTEN protein expression regardless of whether ISG15/ISGylation was knocked out or induced (Fig.S3E), demonstrating that ISGylation in MDA-MB-231 cells does not affect PTEN protein stability. Furthermore, suppression of PTEN would be expected to increase basal pAkt as well as augmenting immediate and peak pAkt. Neither of which we observed (Fig. 3A S.3C and D). Knockout of ISGylation did suppress pAkt activity, but only after 10 minutes of EGF stimulation. This delayed regulation led us to hypothesize that ISGylation did not alter the immediate activation of the pathway but either regulated a feedback loop or was required for efficient receptor trafficking.

EGFR trafficking has been shown to regulate Akt signalling, particularly by sustaining signalling intensity (Ye, Zhu et al., 2016). To explore if the reduction in sustained Akt activation was due to changes in trafficking we analysed EGFR recycling using a surface biotinylation assay (Caswell, Chan et al., 2008). Plasma membrane-labelled EGFR entered the endosomal system and the fraction of EGFR recycled back to the plasma membrane was analysed. Reducing ISGylation (crISG15 and crUBC8) significantly decreased EGFR recycling in comparison to control cells, whereas EGFR recycled at a faster rate when ISGylation was enhanced by USP18 knockout (Figure 4A). These data suggest that ISGylation influences the dynamics of EGFR trafficking, rather than signalling per se.

**Fig. 4.**
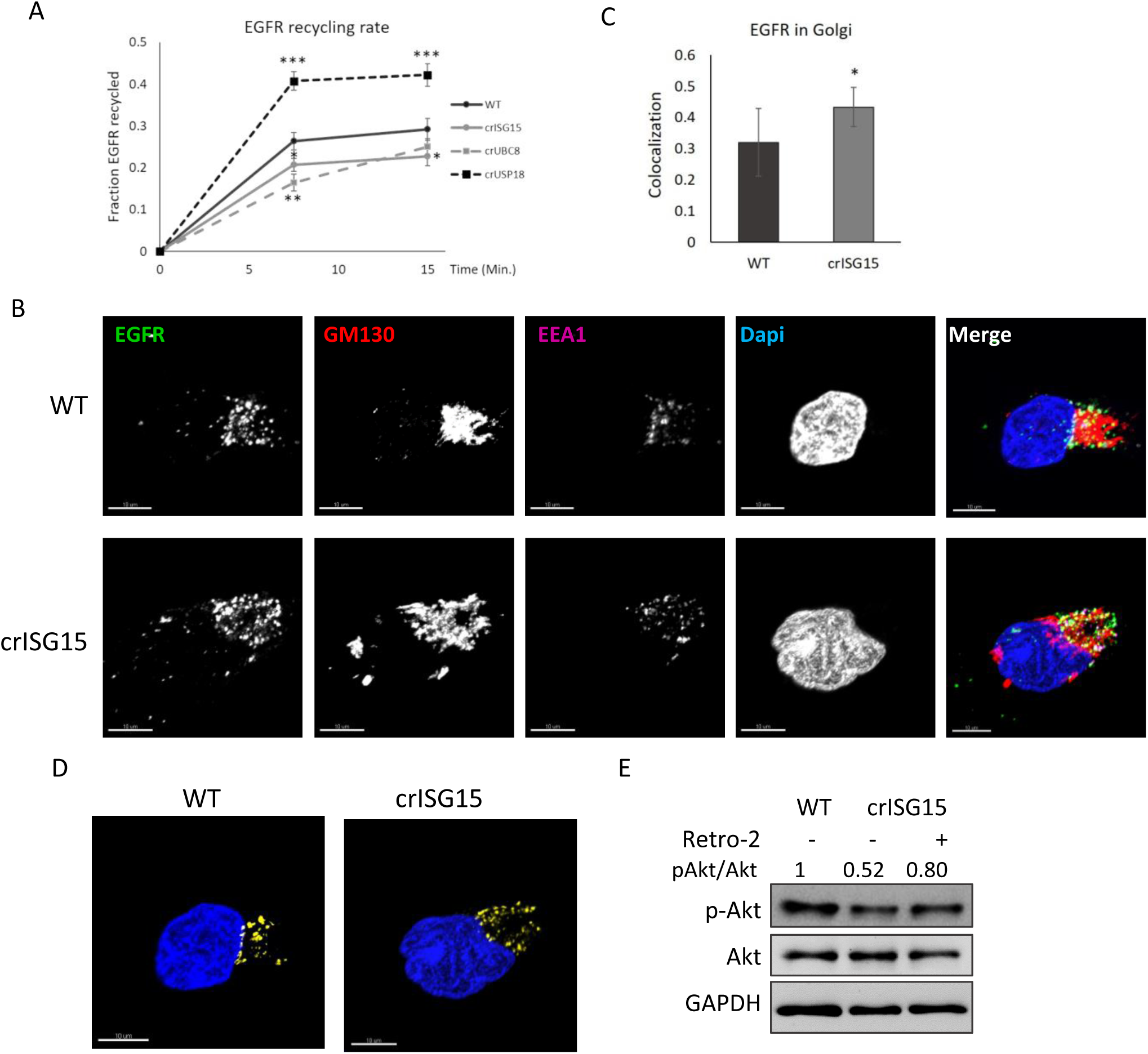
ISGylation promotes faster EGFR recycling. (A) Graph showing EGFR membrane recycling rate following EGF stimulation. The indicated clones were labelled at the membrane with NHS-SS-biotin and then treated with EGF, after the indicated time points, Biotin label was removed from surface proteins at the cell membrane, cells were lysed and the fraction on biotin-labelled-EGFR was analysed. n=5. p-value < 0.05 (*), p-value < 0.01 (**), p-value < 0.005 (***). (B) Representative confocal images, obtained at 60x, of WT and crISG15 cells stimulated for 10 min. with EGF 10ng/ml. Cells were fixed, permeabilised and incubated with antibodies as indicated. Images show EGFR in green, GM130 as Golgi marker in red, the early endosome marker EEA1 in purple and DAPI, used as a reference of the cell location, in blue. 10µm scale bars are displayed in the bottom-left corner. (C) Quantification of EGFR-GM130 co-localization. Bar graph shows the average EGFR fraction which co-localizes with GM130, of the co-localization analysis in WT and crISG15 cells, using Costes method ± SD; n=8. P-value < 0.05 (*). (D) Visualisation of the co-localization between EGFR and the Golgi marker GM130 of the images displayed in (B). In yellow is displayed the fraction of EGFR that co-localizes with GM130 determined using the Imaris software, a colocalization channel was built with it (yellow), DAPI (blue) is shown as a reference. (E) Endosome to Golgi trafficking inhibition induces pAkt levels in crISG15 cells. Blot shows Akt activation of 10min EGF stimulated vehicle-treated WT and crISG15 cells, and crISG15 cells treated with retro-2 for 15min. Quantification of Akt activation, measured as pAkt/Akt signal ratio,normalized to WT one, is showed up to blot panels.

Early endosomes are at the crossroads of EGFR sorting for endocytic recycling, retrograde transport, or degradation through late endosomes/lysosomes. As reduced pAkt levels were not correlated with decreased EGFR levels following EGF stimulation (Fig.3C), we ruled out that ISGylation inhibited the flux of EGFR towards degradation. This indicated that the reduced recycling-rate in the absence of ISG15 was due to inefficient fast recycling or to increase shunting of the receptor toward an alternative route. To test this we analysed EGFR localisation in the context of endosomal trafficking at the time point when we were able to detect ISGylation-dependent changes of Akt activation. Confocal imaging indicated that at 10 minutes EGF stimulation in WT cells the majority of the internalized EGFR co-localized with EEA1-positive endosomal structures, however a small fraction co-localized with GM130 (Fig.4B), a Golgi apparatus marker. In crISG15 cells, we detected a statistically significant increase in the co-localisation between EGFR and GM130 (Fig.4C, D), suggesting that ISGylation plays a role in regulating the trafficking route taken by EGFR through the endosomal system. Furthermore, we found that EEA1, EGFR and GM130 co-localised, which might indicate that EEA1 positive endosomes were fusing to the Golgi, implying increased retrograde transport. To test if retrograde transport reduced Akt activation, we treated the cells with Retro-2, an inhibitor of the endosomes to Golgi transport (Sivan, Weisberg et al., 2016). Treatment with Retro-2 partially rescued pAkt levels in crISG15 cells (Fig.4E), suggesting that reducing EGFR trafficking to the Golgi in crISG15 cells rescues Akt activation.

These data suggest that ISGylation controls regulators of endosomal/retrograde trafficking processes. ISGylation-enhanced EGFR recycling reduces the proportion of the receptor trafficked to the Golgi, increase the residence time in endosomes and returns the receptor to the plasma membrane.

### ISGylation does not have a direct effect on protein stability

To identify the ISG15 substrate/s that control/s endosomal trafficking we devised two complimentary strategies. Firstly, ISG15 has been shown to regulate protein stability (Burks et al., 2014, Desai et al., 2006, Huang et al., 2014), thus we hypothesised that ISG15 could alter signalling by changing expression levels of proteins controlling endosomal trafficking. To test this hypothesis, we devised an expression proteomics screen to assay how inhibiting or enhancing ISGylation alters protein expression. Analysis of protein expression of the different knockout lines showed that 96 proteins were differentially expressed in the clones, when compared to WT cells (table S.1). In addition, we detected three proteins associated with protein trafficking with altered expression levels, the Beta subunit of Sec61 complex (SEC61B); nucleoporin SEH1, or SEHIL, a component of the Nup107-160, subcomplex of the nuclear pore complex; and VAC14 homolog (VAC14), a protein associated with the regulation of phosphatidylinositol 3,5-bisphosphate levels in endosomal membranes. However, all three proteins had the same expression level changes across crISG15, crUBC8 and crUSP18, making them unlikely to be responsible for the opposing changes in EGFR trafficking observed in crISG15/crUBC8 and crUSP18 respectively. Having failed to identify a strong candidate in this initial screen, we devised a more targeted, secondary screen.

We decided to map proteins covalently modified by ISG15 using a direct approach. To determine the ISGylome, we immunoprecipitated endogenous ISG15 from the panel of CRISPR clones, determined which proteins co-immunoprecipitated with ISG15 and used crISG15 as negative control for ISG15 and crUSP8 as negative control for ISGylation. To prevent any non-covalent interaction masking proteins covalently modified by ISG15, we lysed the cells under denaturing conditions prior to the immunoprecipitation step. Analysis of the data revealed 156 proteins as possible ISGylation targets, (Fig.S.4; table S.2) When compared with previous reported ISG15 interactors in the BIOGRID database, 92 were novel targets for ISGylation. Clustering of potential ISGylation targets using STRING (www.string-db.org) (table S.2) indicated that ISG15 is conjugated to proteins associated with a broad range of cell functions, including endosomal trafficking.

Surprisingly, when comparing the results from both screens, only one protein was in common (Fig.S.4). This suggests that: firstly, the effect ISGylation has on Akt signalling or endosomal trafficking is not related to protein stability, and secondly that ISGylation per se does not primarily regulate protein stability under basal conditions.

### ISGylation reduces GDI2 affinity for Rabs

One of the putative ISGylation substrates reported to regulate endosomal trafficking was GDP Dissociation Inhibitor 2, also known as RabGDIB or GDI2 (Fig.5A). GDI2, a member of the GDI family of proteins, is a ubiquitously expressed cytoplasmic protein that interacts with members of the small GTPases Rab protein family (Rab). GDIs bind Rabs in their inactive, GDP-bound state. The mechanism involves initially a protein-protein interaction between the Rab and the GDI and then a more stable interaction between the GDI lipid binding domain and Rab C-terminal prenyl group (Ignatev, Kravchenko et al., 2008). GDI2 is a regulator of Rab activity and localisation making it a plausible candidate to be an integrator of ISGylation and EGFR endosomal trafficking. Initially, we decided to confirm that GDI2 is indeed ISGylated. We overexpressed ISG15 and GDI2 in WT as well as in crUBC8 cells as negative controls. We transfected Myc-DDK tagged GDI2, Strep-tactin tagged ISG15 or both, and determined the presence or absence of ISGylated GDI2 by performing a Strep-tactin-ISG15 pull-down under denaturing conditions. We detected the presence of exogenous GDI2 with a shift of the apparent molecular weight only in WT cells transfected with both Myc-DDK-GDI2 and Strep-tactin-ISG15 (Fig.5B). The absence of GDI2 in crUBC8 and in the additional negative controls confirmed our hypothesis that GDI2 is ISGylated under basal conditions.

**Fig. 5.**
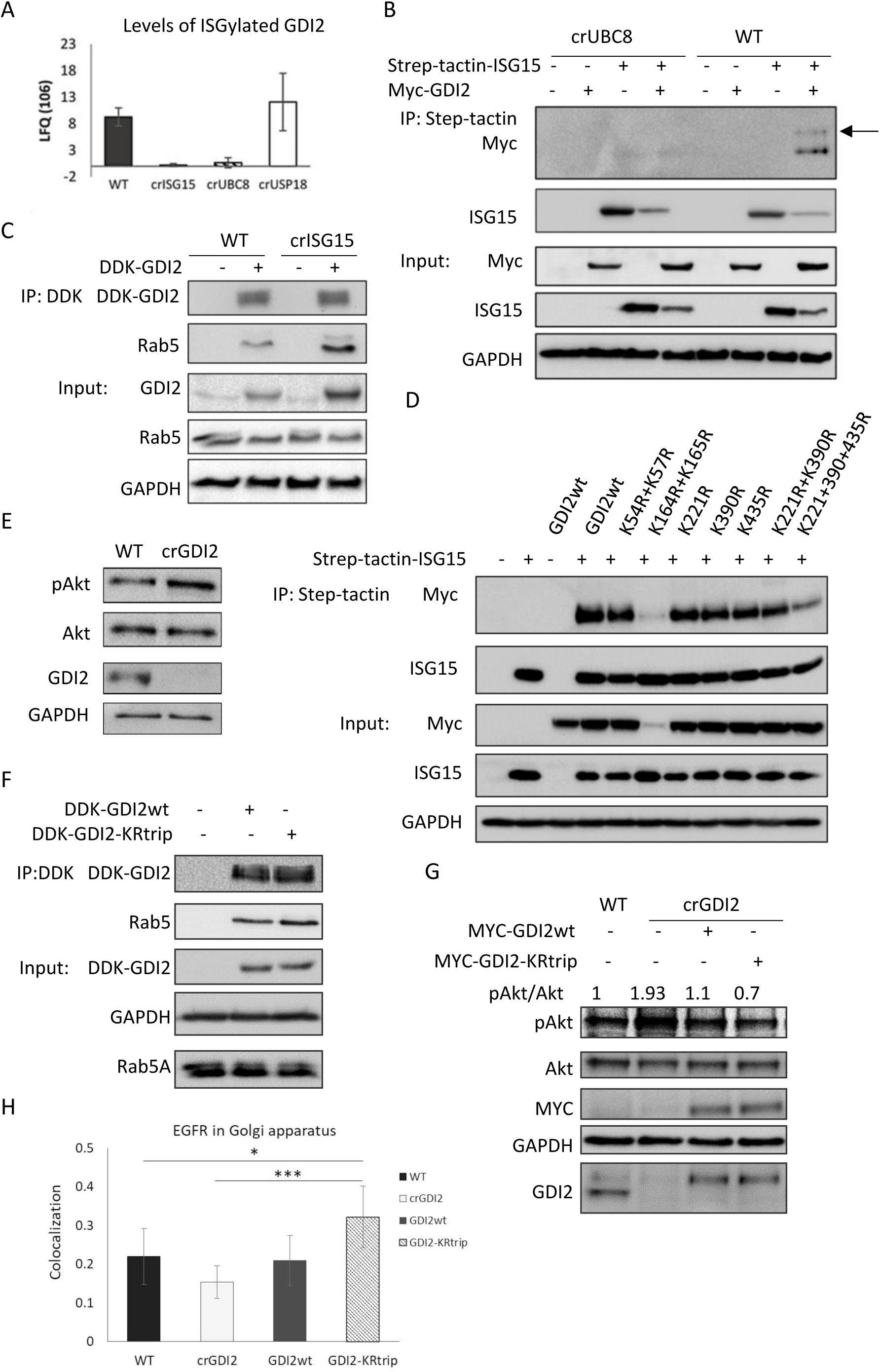
ISGylation of GDI2 reduces its activity and increases Akt activation. (A) LFQ values for GDI2 obtained from ISG15 pull-downs of the indicated clones, analysed by mass spectrometry, protein identification and quantification was performed using MaxQuant. (B) Analysis of GDI2 ISGylation using crUBC8 and WT cells transfected with empty vector, Step-tactin-ISG15, MYC-DDK-GDI2 or both, and subjected to a Step-tactin pulldown. Blots show the pulldown and a 5% of the total lysates (Input). Arrow indicates ISGylated GDI2. (C) WB of WT and crISG15 cells transfected with MYC-DDK-GDI2, after 48h cells were lysed and subjected to an anti-Flag pull-down, GDI2 activity was analysed by the levels of Rab5 detected in the pulldowns. Blots show the pulldowns and a 5% of the total lysates (Input). (D) Analysis of GDI2 ISGylation sites. WT cells were transfected with Strep-tactin ISG15, MYC-DDK-GDI2, or Strep-tactin ISG15 with either MYC-DDK-GDI2wt or the indicated GDI2 mutants. After 48h, cells were lysed and subjected to a Strep-tactin pull-down. The ISGylation status of the different GDI2 constructs was measured by the levels of GDI2 detected. Blot shows the results of the pulldown and a 5% of the total lysates (Input). (E) Analysis of Akt activation in crGDI2 by WB of WT and crGDI2 cells stimulated with EGF 10ng/ml for 10min. (F) WB of WT cells transfected with MYC-DDK-GDI2wt, or MYC-DDK-GDI2-KRtrip and after 48 h were lysed and subjected to Flag IP. GDI2 activity was measured by the levels of Rab5 detected in the pulldowns. Blots show the pulldowns and a 5% of the total lysates (Input). (G) Analysis of the effect of GDI2-KRtrip in Akt activation by WB of lysates from WT cells, crGDI2 cells, crGDI2 cells transfected GDI2wt or transfected with the GDI2-KRtrip expression vector for 48h, treated for 10min. with EGF. Values at in the upper part of the WB show the pAkt/Akt ratio as measure of Akt activation, normalized to the WT cells value. (H) Quantification of EGFR-GM130 co-localization. Bar graph of the average EGFR fraction that co-localizes with GM130 using Costes method ± SD of WT cells, crGDI2 cells, crGDI2 transfected with GDI2 expression vector, GDI2wt, or with the mutant GDI2, GDI2-KRtrip, treated for 10min. with EGF. n=8. p-value < 0.05 (*), p-value < 0.01 (**), p-value < 0.005 (***).

ISG15 has a broadly described role as protein stability regulator, however neither the protein expression screen (Fig. S5A) nor a targeted western blot analysis of GDI2 levels (Fig.S5B) showed any changes in GDI2 protein levels in the different cell lines. Therefore, we hypothesized that ISGylation of GDI2 may be affecting GDI2 function rather than its stability. To test this hypothesis, we analysed the ability of GDI2 to interact with Rabs in the presence or absence of cellular ISGylation. We transfected Myc-DDK-GDI2 into WT, crISG15, crUBC8, crUSP18 and analysed the interactome in an unbiased manner by mass spectrometry (Fig.S5C; tableS.3). We found several Rabs to be interacting specifically with GDI2. Intriguingly, both Rab5 and Rab11 co-precipitated to higher levels with GDI2 in the samples derived from crISG15 and crUBC8. Suggesting that ISGylation limits the formation of a GDI2-Rab5/11 complex. In contrast, association of Rabs with GDI2 decreased further in crUSP18 cells. To corroborate these results independently we transfected WT and crISG15 cells with Myc-DDK-GDI2 and assayed endogenous Rab5 in the immunoprecipitate by Western blotting (Fig.5C). Confirming our mass spectrometry data, GDI2 co-precipitated higher levels of Rab5 in the crISG15 when compared to WT. Surprisingly, transfected exogenous GDI2 was expressed at consistently increased levels in crISG15 cells (Table S.3). However, this effect was absent in crUBC8, which indicates that this is not dependent on ISGylation. Rab5 and Rab11 are required for the maturation of early and recycling endosomes respectively (Gorvel, Chavrier et al., 1991, Wilcke, Johannes et al., 2000). Detecting that ISGylation alters the complex formation between a GDI2 and these two Rabs suggests that regulation of the interaction between GDI2 and RabGTPases, and consequent association of active Rab5/11 with endosomal membranes, may be responsible for the observed deregulation of EGFR recycling and downstream signalling.

### ISG15 modified several lysine-residues of GDI2

To identify potential ISGylation sites on GDI2 we overexpressed GDI2 in COS1 cells, and treated the cells with INFb1 to boost endogenous ISGylation levels. We lysed the cells in presence of protease/DUB inhibitors, affinity purified GDI2 and digested the protein with trypsin. Analysis by LC-MS/MS indicated that lysine 435 (Fig.S5D and S5F) was modified by a double glycine peptide, which is a residual tag consistent with a modification by several Ubiquitin-like proteins, including ISG15. INFb1 treatment increased the level of the modification, suggesting that this may be an ISGylated residue. In addition to this unbiased approach, we mined databases for potential ISGylation sites. ISG15 and ubiquitin share the same Lysine-Glycine-Glycine (K-GG) peptide mark when digested with trypsin and are therefore undistinguishable from each other in standard ubiquitination screens, where anti-K-GG antibodies are used to enrich modified peptides. We interrogated the Phosphosite database (www.phosphosite.org) for K-GG sites detected on GDI2 and found that peptides containing lysine 54, 57, 164, 165, 221 and 390 have been identified with the Ubiquitination/ISGylation marker.

To test if any of these sites could be ISGylated we substituted each lysine for arginine, creating the following mutants: K45R/K57R, K164R/K165R, K221R, K390R, K435R as well as combinations of several locations such as K221R/K390R and K221RK390R/K435R. Analysis by Strep-tactin-ISG15 pull-downs of the different GDI2 constructs (Fig.5D) showed that no single site mutant significantly decreases ISGylation. The double mutant, K164R/K165R, showed a marked decrease in ISGylation, but this was linked to a decrease in protein level, which suggest that those mutations were detrimental for protein stability. A marked reduction in ISG15 conjugation was detected in the GDI2 with triple mutation, in K 221, 390 and 435, which suggest that ISG15 can be conjugated to multiple sites.

We had observed that GDI2 binding to Rabs is enhanced in cells devoid of ISGylation (Fig. 5C and S5C) and that the absence of ISGylation coincided with an impairment of Akt activation upon mitogen stimulation (Figs. 3A and B). To investigate if those two events are causally linked, we knocked-out GDI2 in MDA-MB-231-luc-D3H2LN cells (crGDI2). Firstly, to determine if GDI2 is regulating Akt activation, we treated WT and crGDI2 cells with EGF for 10min. crGDI2 cells responded with a two-fold higher pAkt level compared to control cells upon EGF stimulation (Figs.5E, S5E), suggesting that knockout of GDI2 enhances signalling through the PI3K/Akt pathway. Secondly, our data suggested that ISGylation of GDI2 on specific residues inhibits the ability of GDI2 to interact with Rabs. To test this hypothesis we transfected WT cells with either GDI2wt or the triple mutant K221RK390R/K435R (GDI2-KRtrip), lysed the cells and immunoprecipitated GDI2 and assessed the ability of GDI2wt and GDI-KRtrip to interact with Rab5. We detected a higher binding of Rab5 to GDI2-KRtrip (Fig.5F), confirming that reducing ISGylation of GDI2 increased the ability of GDI2 to interact with Rabs. Thirdly, to tie these results together and to exclude the possibility that ISGylation and loss of GDI2 are independently linked to pAkt signalling, we performed a rescue experiment. We rescued crGDI2 cells by transfecting them with either GDI2wt or GDI2-KRtrip and analyse their ability to reverse the increase of pAkt. Reassuringly, re-expression of GDI2wt was able to rescue and reduce pAkt to a level similar to the WT cells (Fig.5G). More interestingly, GDI2-KRtrip reduced pAktser levels even further (Fig.5G).

The partial rescue of pAkt levels in crISG15 treated with Retro-2 (Fig. 4E) suggested that the retrograde transport to the Golgi is associated with the reduced Akt activation in ISGylation deficient cells. To determine if the enhanced activation of Akt caused by GDI2 knockout is associated with reduced EGFR trafficking to the Golgi, we analysed EGFR co-localising with the Golgi, after 10min. of EGF simulation in WT cells, crGDI2 cells and in crGDI2 rescued by expressing GDI2wt or GDI2-KRtrip. When quantifying co-localisation of EGFR with a Golgi marker by confocal microscopy we found that knocking out GDI2 reduced the presence of EGFR at the Golgi when compared to WT cells. This reduction was rescued when GDI2 was re-expressed. Intriguingly, rescuing crGDI2 with GDI2-KRtrip further increased the localisation of EGFR to the Golgi to a level beyond the WT cells (Figs.5H and S6A). EGFR was co-localising with the majority of the Golgi indicating that, in these cells, the endosomes had fused with the Golgi (Fig.S6B).

Taken together, these experiments demonstrate that GDI2 is ISGylated, ISGylation of GDI2 inhibits binding to Rabs and inhibition of GDI2 ISGylation suppresses Akt signalling by controlling the routing of EGFR through endosomes and the Golgi.

### ISG15 expression correlates with Akt-signalling in human tumours

Our *in vitro* data showed that in a basal-like cell line ISGylation enhanced sustained Akt signalling (Fig.3A and 3B). To determine if the correlation between ISG15/ISGylation and Akt signalling was also observable in human breast tumours, we mined The Cancer Genome Atlas (TCGA) database (http://cancergenome.nih.gov/).TCGA has no protein expression data for ISG15 or UBC8, for this reason we used mRNA levels as surrogates. Initially, to establish if we could assume that high levels of ISG15 would correlate with high levels of ISGylation, we determined if there was correlation between ISG15 levels and UBC8, the E2 ligase. Analysis of mRNA expression levels in primary tumours (Fig. 6A), revealed that these genes were highly correlated (Spearman’s correlation=0.66, p < 2.2e-16), indicating that high ISG15 could be taken as indicative of high ISGylation. With this assumption, we mined the database for a possible correlation between ISG15 mRNA levels and a gene signature which had been associated with high PI3K/Akt pathway activation in human breast cancers (Creighton, 2007). The Analysis revealed that there is a statistical significant correlation between high ISG15 mRNA levels and enhanced Akt pathway activity in basal, luminal A and luminal B subtypes in both, lymph node negative (Fig.6B) and positive status (Fig.6C). In addition to this, we wanted to determine if pAkt levels correlated with ISG15 mRNA levels in breast cancer patient samples. TCGA includes a data set for which pAkt has been measured by RPPA. In this dataset ISG15 mRNA expression showed no correlation with pAkt in lymph node negative patients, but a modest negative one for luminal B (Fig 6D), however in basal tumours with lymph node metastasis a positive correlation between ISG15 mRNA and pAkt was found (Fig.6E) (P-value = 0.0012). Interestingly, the same correlation was also found in for UBE2L6 mRNA levels and p Akt in Luminal B (P-value= 0.0169) and basal tumours (P-value = 0.0182) with lymph node metastasis (Fig.6F). This suggest that in human breast cancer ISGylation correlates with enhanced pAkt and signalling though the PI3K/Akt pathway.

**Fig. 6.**
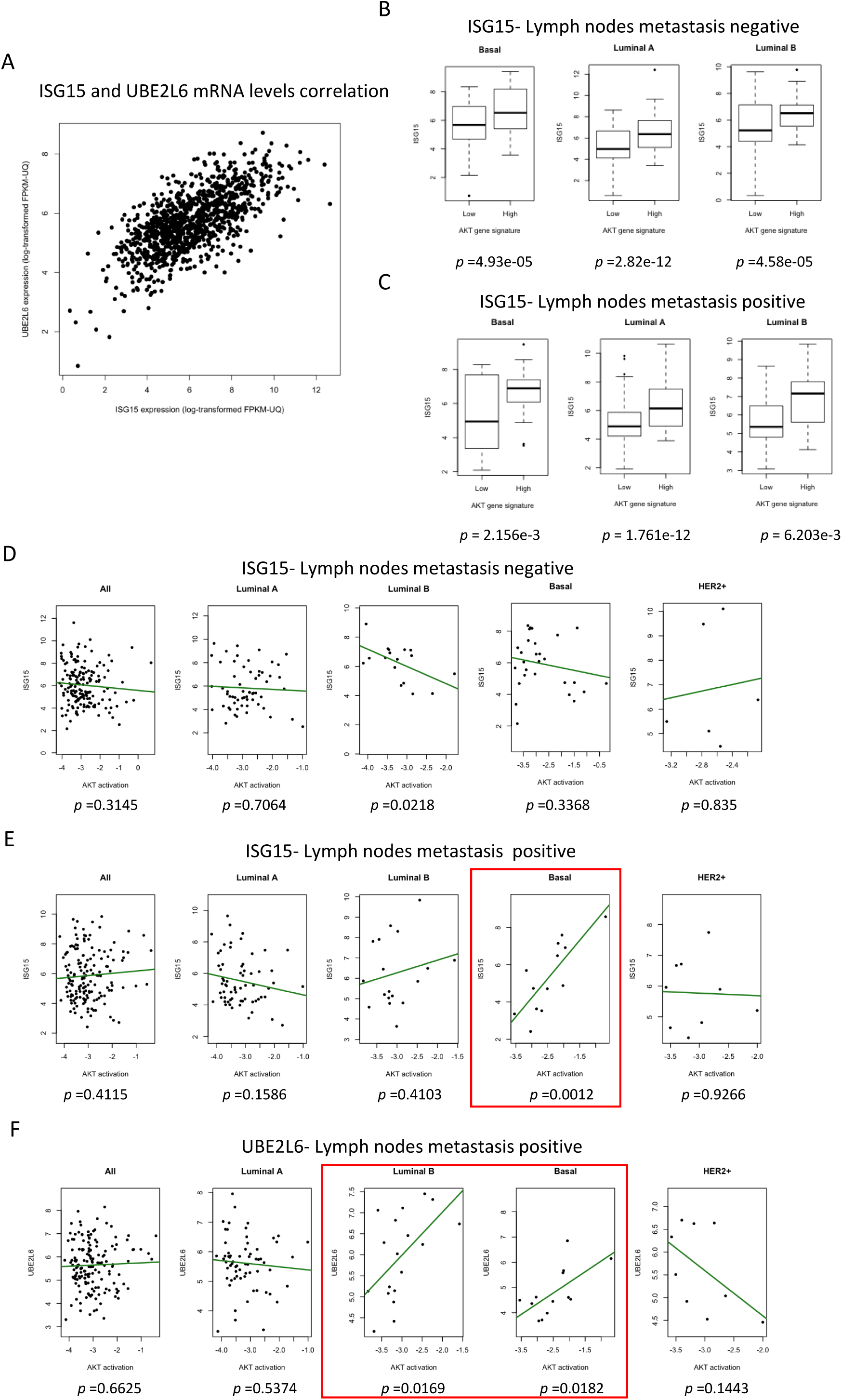
ISG15 positively correlates with Akt activation in breast cancers. (A) Correlation analysis of ISG15 and UBE2L6 mRNA expression levels in breast cancer samples. Spearman correlation value is displayed in the top right. (B) Boxplots showing Akt pathway activity gene signature correlation with mRNA levels of ISG15 in breast cancer in lymph node negative samples for the subtypes basal (n=68), luminal A (n=174), and luminal B (n=41). p-values are displayed below each graph. (C) Boxplots showing Akt pathway activity gene signature correlation with mRNA levels of ISG15 in breast cancer in lymph node metastasis positive samples in the subtypes basal (n=36), luminal A (n=188) and luminal B (n=53). p-values are displayed below each graph. (D) Correlation in tumours of patients with no lymph node metastasis as a whole (n=166), or classified by subtype; luminal A (n=62), luminal B (n=17), basal (n=28) and HER2 (n=6). (E) Correlation in tumours of patients positive for lymph node metastasis as a whole (n=156), or classified by subtype: Luminal A (n=65), Luminal B (n=18), Basal (n=13) and HER2+ (n=9). (F) As (B) for UBE2L6 expression. In red boxes the subtypes that showed statistical correlation.

Taken together, these data supports our *in vitro* data and the hypothesis that high levels of ISGylation enhances the malignancy of breast cancer tumours by promoting Akt signalling. Mechanistically, we propose that ISGylation of GDI2 achieves this. ISGylated GDI2 has a lower affinity for Rab proteins, which in turn induces faster receptor recycling to the membrane by reducing the flux of EGFR towards the Golgi. Faster receptor recycling to the membrane then enables the tumour cells to sustain PI3K/Akt activation.

## Discussion

Metastasis is a main determinant of cancer mortality and an understanding of how the primary tumour acquires the ability to form secondary cancers is key to target the process. Altered EGFR-family RTK signalling is most commonly associated with the induction of aggressive tumour characteristics, such as invasion, proliferation or angiogenesis in breast tumours (Butti, Das et al., 2018, Masuda, Zhang et al., 2012). To complement the picture, our data shows that ISG15 expression and prognosis are inversely associated in patients with lymph nodes metastasis (Fig.1B-C) with the highest association in the Luminal B and basal-like subtype (Fig.S1A). We show that ISGylation enhances EGFR recycling, effectively increasing membrane localisation and signalling. Consistently, EGFR overexpression has been associated with poor prognosis in both these subtypes (Liu, He et al., 2012, Park, Jang et al., 2014). Conversely, the HER2+ subtype shows no correlation between ISG15 and either prognosis (Fig.S1A) or Akt activity (Fig.7), which suggest that HER2 overexpressing cells are indifferent to ISG15-enhanced trafficking. HER2 positive tumours are defective for receptor internalization after ligand stimulation (Hommelgaard, Lerdrup et al., 2004), which would in turn render them resistant to altered downstream trafficking. This may explain why there is no correlation between ISG15, Akt signalling and mortality in HER2-overexpressing tumours.

Consistent with patient data and in agreement with previous studies (Burks et al., 2014, Desai et al., 2012), depleting ISGylation, reduces PI3K/Akt signalling and, consequently, the aggressive potential of tumour cells. The association between ISG15 and Akt has been previously observed in macrophages (Yanguez et al., 2013), where it is linked to phagocytosis. It appears that high levels of ISGylation has allowed tumour cells to hijack an immune cell mechanism to their own benefit.

Early endosomes are a platform for the PI3K/Akt/GSK3b signalling axis (Schenck, Goto-Silva et al., 2008). Several studies have shown that increased localisation of EGFR in early endosomes increases the metastatic potential by enhancing signalling through the PI3K/Akt axis (Caswell et al., 2008, de Graauw, Cao et al., 2014, Nishimura, Takiguchi et al., 2015). In this context it is unsurprising that expression of other regulator of EGFR endocytosis, such as Rab5, are markers of poor prognosis (Frittoli, Palamidessi et al., 2014) and lymph node metastasis in breast cancers (Yang, Yin et al., 2011).

We show that ISGylation levels also directly correlate with high recycling rates and decreased flux towards the Golgi. This effect is due to the ISGylation of GDI2, a regulator of Rab localisation and activity (Shisheva, Chinni et al., 1999), GDI2 contains two conserved domains, a protein-protein interaction domain, the Rab-binding platform, and a protein-lipid interaction domain, or lipid binding pocket, connected by the GDI effector loop (Luan, Heine et al., 2000). Of the three sites identified as ISGylated, lysine 221 may be the most relevant for the Rab interaction. Sequence alignments suggest that it is at the junction between the hinge and the lipid-binding domain, towards the side of the protein that interacts with Rabs. It is plausible that ISGylation of Lysine 221 could limit the interaction with the C-terminal prenyl group of the interacting Rab, the key step for the GDI-mediated extraction of Rab-GTPases from the membrane (Gilbert & Burd, 2001). Lysine 390 and Lysine 435 on the other hand, are probably facing away from the Rab-interaction domain. It is therefore unlikely they would directly regulate the Rab-GDI2 interaction. Further research is needed to establish how ISGylation of these residues regulates GDI2 activity, nevertheless it is likely that ISGylation limits the ability of GDI2 to retrieve and/or retain inactive RabGTPases. In this context changes in Rab5 and Rab11 availability could induce a rerouting EGFR away from a retrograde route and support Akt activation, either by decreasing the time in the Trans-Golgi network compared with the endosomal one or by increasing the levels of EGFR in different types of endosomal compartments. It is tempting to hypothesize that receptor trafficking is dependent on the balance between Rabs and GDIs, thus in different cell types with different expression levels, modification of GDI2 availability for Rab-binding by ISGylation may effect RTKs mediated signalling distinctly.

In conclusion, we have identified a novel pathway involving ISG15, GDI2, Rabs, which potentiates breast cancer progression by shunting EGFR signalling towards sustained PI3K/Akt signalling.

## Acknowledgements

This work was supported by Breast Cancer NOW (2013NovPR183), Cancer Research UK. Cancer Research UK (CRUK Edinburgh Centre C157/A255140), Wellcome Trust (Multiuser Equipment Grant, 208402/Z/17/Z).

## Experimental Procedures

### Cell lines

MDA-MB-231 subclone D3H2LN, Cos1 and HEK293t were gown in DMEM 4.5 g/l glucose supplemented with 10% foetal bovine serum and 2mM glutamine, at 37°C and 5% CO2.

### Reagents

Recombinant Human Interferon Beta-1a was obtained from Prospec; EGF, insulin, ampicillin, kanamycin and puromycin were obtained from Sigma; transfection reagent TransIT-X2® was obtained from Mirus; transfection reagent jetPRIME was obtained from Polyplus transfections.

Plasmids, cloning and GDI2 mutagenesis: Knock-out cell lines were created using two different gRNAs sequences against the different genes, ISG15 (crISG15-1 Fw 5’ GCTGGCGGGCAACGAATTCC 3’ and crISG15-2 Fw 5’ CTGCGTCAGCCGTACCTCGT 3’), UBE2L6 (crUBC8-1 Fw 5’ CTGTCCGTTCTCGTCCACGT 3’ and crUBC8-2 Fw 5’ GGCTTGAACGGATACTCCGG 3’) and USP18 (crUSP18-1 Fw 5’ TCACGAATGAGCAAGGCGTT 3’ and crUSP18-2 Fw 5’ GCAAATCTGTCAGTCCATCC 3’), GDI2 (crGDI2-1 Fw 5’ GCCACCCGAGTCAATGGGGA 3’ and crGDI2-2 Fw 5’ CACTCTCTCCTCCGTACGTA 3’) into a lentiviral CAS9 expressing plasmid (Addgene Plasmid # 49535) using the BSMB1 restriction sites. gRNAs for ISG15, UBE2L6 and USP18 were obtained from (Shalem, Sanjana et al., 2014). Plasmids were transformed into Stbl3™ E. coli strain (Thermo Fisher). Lentiviral particles were synthetized through transfection of 20µg of CAS9 expression vector, 2µg of the envelope plasmid pCMV-pVSVg (AddGene Plasmid #8454) and 7.5µg of the packaging plasmid psPAX2 (AddGene Plasmid #12260) in HEK293t cells. Media was recovered every 24h for 3 days, centrifuged at 1500rpm for 3 minutes and filtrated through a 0.45µm filter prior to be added to MDA-MB-231 cells. After 48h cells were selected to puromycin selection (2µg/ml), individual clones were grown and tested for protein expression. Two clones per guide were used in the experiments, treated as biological replicate. Transfection were carried out using transit system, using a relationship of 2µl of transfection reagent was used per microgram of plasmid in DMEM-serum free medium.

Human ISG15 expressing plasmid was obtained by cloning ISG15 cDNA from MDA-MB-231 RNA, using the Forward primer 5’ATGGGCTGGGACCTGACGGTG 3’ and the reverse primer 5’ TTAGCTCCGCCCGCCAGGC 3’. PCR product was gel purified using the QuiaQUICK gel extraction kit and cloned into the pCR8/GW/TOPOR entry vector. Plasmids were sequenced to confirm the presence and orientation of the inset by using the GW1 and GW2 primers included in the kit. Once characterized, one clone was transferred into a puromycin-resistant derivative of TAPE5-N destination vector 1 using LR Gateway Technology.

Human GDI2 expression vector pCMV6-DDK-Myc-GDI2, was bought from Origene (cat. RC200596). GDI2 mutagenesis was carried out using the Q5® Site-Directed Mutagenesis Kit (Promega, cat. E0554S) using manufacturer recommendations. The following primers for site-directed mutagenesis were designed using the online tool provide by New England Biolabs (url://nebasechanger.neb.com/): For K54+54R (Fw 5’ TTTAGAATACCAGGATCACCACCC 3’ and Re 5’ TCTTCTGTATAAATCTTCCAATGGTGTTATAG 3’); K164+165R (Fw 5’ ATTGATCCTAGGAGGACCACAATGCGAGATGTGTATAAGAAATTTGAT 3’ and Re 5’ TGTGGTCCTCCTAGGATCAATGCCTTCAAAAGTTCTTGGATCTT 3’); for K221R (Fw 5’ AAGATATGGCAGAAGCCCATACC 3’ and Re 5’ GCCAAAGATTCACTGTAAAG 3’); for K390R (Fw 5’ CCTGGTACCAAGAGACTTGGGAA 3’ and Re 5’ AGGTCACTGATGCTAACAAATTTC 3’); for K435R (Fw 5’ TGAGGAAATGAGGCGCAAGAAGA 3’ and Re 5’ AAGTCAAACTCTGATCCTGTC 3’). Phenotype assessment: Proliferation assays were performed by plating 25000 cells of each construction plates and counting the amount of cells every 12h for 3 days. Proliferation rate was determined as the slope of the variation in cell number vs time.

Soft agar assays were performed as 3 T-6 well plates were coated with 1ml of 0.6% agar in DMEM and keep at room temperature until solidified, then 1ml of 0.3% agar in DMEM with 1 10^4^ cells were applied to the top, once solidified 1ml of media was applied on top. Media was changed twice per week, after 4 weeks colonies were detected using crystal violet staining.

Migration and invasion assays, were performed using the IncuCyte ZOOM system (Essen Biosciences), following manufacturer protocol for scratch wound experiments in matrigel. 5 10^4^ cells were plated per well in a 96-well plate and a scratch-wound was made. Phase-contrast images were taken every 3h, at 10x. Invasion and motility were measured as percentage of wound closed per time point using IncuCyte ZOOM analysis software.

### Immunoblots

For protein and phosphoprotein expression analysis cell were lysed in lysis buffer (0.1% triton x-100, 50 mM HEPES pH 7.5, 150 mM sodium chloride, 1.5 mM magnesium chloride, 1 mM EGTA, 1 mM sodium fluoride, 10mM Beta-glycerophosphate, 1 mM sodium vanadate, 10 µM leupeptin, 100 nM aprotinin, 1 mM PMSF). Cleared lysates were resolved using 10% or 12% acrylamide gels and proteins were transferred to PVDF membranes (Whatman) using Mini Trans-Blot® Electrophoretic Transfer Cell (BIO RAD). Membranes were blocked in 4% BSA, incubated in primary antibodies (see paragraph below); secondary HRP antibodies, from Cell Signalling, were used at 1:10000 dilution. Membranes were visualized using ClarityTM Western ECL Substrate (BioRAD) in a ChemidocTM MP (BioRAD). Western blot quantification were performed using ImageJ.

Anti-ISG15 antibody (1:1000) was bought from PBL Assay Science; antibodies against GDI2 (1:2000) were bought from Proteintech and Life Technologies; antibody against USP18 (1:500) was bought to Proteintech; antibodies anti pEGFR Tyr1086 (1:2000), pAkt Ser473 (D9E) (1:2000), Akt (1:1000), pGSK3b Ser9 (S9) (1:2000), GSK3b (27C10) (1:2000), pS6K1 Thr421/Ser424 (1:1000), S6K1 (49D7) (1:1000), MYC-tag (71D10) (1:1000), Rab5 (C8B1) (1:1000), PTEN (D4.3) (1:3000) and GAPDH (14C10) (1:3000) were obtained from CST; antibody against pAkt Thr308 (1:500) was obtained from Millipore; antibodies against HRP-FLAG® (1:3000), ppERK Thr202/Thr185,Tyr204/Tyr187 (1:3000) and ERK (1:3000) were bought from Sigma; antibodies against UBC8 (K1H3) (1:500) and EGFR (C-30) (1:3000) were obtained from Santa-Cruz.

Reverse phase protein array (RPPA): Biological triplicates of each construction were lysed in lysis buffer (1% triton x-100, 50 mM HEPES (pH 7.5), 150 mM sodium chloride, 1.5 mM magnesium chloride, 1 mM EGTA, 100 mM sodium fluoride, 10 mM sodium pyrophosphate, 1 mM sodium vanadate, 10% glycerol, 10 µM leupeptin, 100 nM aprotinin). Cleared lysates were normalized to 1.5mg/ml and four serial dilutions of each sample, were spotted onto nitrocellulose-coated slides (Grace Bio-Labs) in technical duplicates. Primary antibodies were applied at 1:250 concentration. Bound antibodies were detected by incubation with DyLight 800-conjugated secondary antibody (New England BioLabs). An InnoScan 710-IR scanner (Innopsys) was used to read the slides, and images were acquired at the highest gain without saturation of the fluorescence signal. The relative fluorescence intensity of each sample spot was quantified using Mapix software (Innopsys). The signal intensity of each antibody was determined using the linear regression made with the median intensity of the four different sample dilutions. Signal intensity were normalised to total protein loaded, determined by staining a slide with fast-green, a non-specific protein dye and scanning at 790Um.

Analysis of changes in whole-proteome and ISGylome by Liquid Chromatography-Tandem Mass Spectrometry: Changes in whole proteome levels was analysed with the FASP-protocol (Wisniewski, Zougman et al., 2009). 150mm plates were lysed in lysis buffer supplemented with 1% SDS, sonicated, and protein concentration was determined by BCA. 100 micrograms of protein was used per clone. Proteins were isolated and subjected to Lys-C and trypsin digestion. Peptides concentration were measured using a Nanodrop ND-1000.

10µg of peptides were purified using the in-stage tips protocol 5. The eluted peptides were lyophilized in a Concentrator Plus (Eppendorf), resuspended in 0.1% TFA and analysed by LC-MS/MS on a QExactive™ Hybrid Quadrupole-Orbitrap™ Mass Spectrometer (Thermo Fisher) as described (Turriziani, Garcia-Munoz et al., 2014). Protein identification and quantification was performed by label-free quantification using MaxQuant software (Cox & Mann, 2008).

### ISG15 Pulldowns

For ISGylome analysis, 10cm dishes per experiment with two different clones per knock-out, were grown. Cells were lysed in lysis buffer with 1% SDS and diluting the cleared lysate by a 10 fold. Pulldowns were performed in a KingFisher Duo (Thermo Scientific). Cleared lysates were incubated with protein G Mag Sepharose™ Xtra beads (GE Healthcare) and 1µg of antibody anti-ISG15 per sample for 4h, washed two times in lysis buffer and three times in TBS and resuspended in digestion buffer (2M urea, 50Mm Tris-HCL pH7.5, 1 mM DTT) with 1µg of porcine trypsin MS-grade (Promega) for digestion. Peptides were desalted and purified using STAGE tips. The eluted peptides were lyophilized in a Concentrator Plus (Eppendorf), resuspended in 0.1% TFA and analysed by LC-MS/MS on a QExactive™ Hybrid Quadrupole-Orbitrap™ Mass Spectrometer (Thermo Fisher) as described in (Turriziani et al., 2014). Protein identification and quantification was performed by label-free quantification using MaxQuant software (Cox & Mann, 2008).

For exogenous ISGylation analysis cleared lysates were incubated with strep-tactin MagStrep “type3” XT beads (IBA Biosciences) for 2h, washed two times in lysis buffer and one time in TBS and resuspended in loading buffer without reduction agents.

### GDI2 Pulldowns

10cm dishes Cell were lysed in Lysis buffer (see Immunoblotting section above); cleared lysates were incubated with Anti-FLAG® M2 Magnetic Beads (SIGMA) for 2h and washed two times in lysis buffer and one time in TBS. For Immunoblot analysis beads were resuspended in loading buffer supplemented with 1mM DTT. For MS analysis, beads were washed two extra times in TBS and resuspended in digestion buffer (2M urea, 50Mm Tris-HCL pH7.5, 1 mM DTT) with 1µg of porcine trypsin MS-grade (Promega) for digestion. Peptides were desalted and purified using the STAGE tips. The eluted peptides were lyophilized in a Concentrator Plus (Eppendorf), resuspended in 0.1% TFA and analysed by LC-MS/MS on a QExactive™ Hybrid Quadrupole-Orbitrap™ Mass Spectrometer or Orbitrap Fusion™ Lumos™ Tribrid™ (Thermo Fisher) as described (Turriziani et al., 2014). Protein identification and quantification was performed by label-free quantification using MaxQuant software (Cox & Mann, 2008).

### Imaging

Cells were seeded onto a glass coverslip, starved overnight and treated for 10min. with EGF (10ng/ml). Cells were washed twice with PBS, fixed and permeabilised with 3.7% formaldehyde, 0.1% NP-40 in 50mM Pipes pH6.8, 125mM NaCl, 10mM MgCl2, 10mM EGTA, for 5min. and blocked in TBS 2% BSA for 1h. Coverslips were incubated overnight with rabbit anti EGFR-Alexa488 (Abcam), 1:100, rabbit anti GM130 -Alexa647, 1:100 (Thermo Scientific) and mouse anti EEA1 (BD Biosciences), 1:200, overnight. Slides were washed twice with TBS and then incubated for 1 h at room temperature with secondary antibody anti-mouse Alexa-546 (Life Technologies). Slides were washed twice with TBS and incubated with DAPI, 1:100, for 5 minutes, washed two times and mounted using VECTASHIELD antifade mounting media (Vector labs). Stack images were taken with an Olympus FV100 confocal, with 60x oil objective. Co-localization analysis was performed using Imaris 7.7 software, images were subjected to background subtraction and co-localization was determined using the Costes method included in the ImarisColoc package.

### EGFR recycling

Recycling assays were performed as described previously (Caswell et al., 2008). Briefly, cells were labelled at 4°C with 0.13mg/ml NHS-SS-Biotin (Pierce), and biotinylated receptors internalized for 30 min at 37°C in serum free medium before removal of remaining surface biotin by reduction with sodium 2-mercaptoethane sulfonate. Internalised receptors were chased from the cells by returning the cells to 37°C, surface biotin removed by a second reduction step and the fraction of biotinylated integrins analysed by ELISA. The antibodies used for capture ELISA detection of EGFR were obtained from BD Biosciences.

Statistical analysis of Akt signalling and ISG15 expression in human breast tumours Data acquisition and pre-processing: Publicly available RNA-seq data, in FPKM format, from breast tumours in the TCGA-BRCA project, was downloaded from the genomic data commons (https://portal.gdc.cancer.gov/). Non-expressing genes were excluded and FPKM counts were upper quartile normalised, scaled and log2(x+1) transformed. Breast cancer subtypes were defined using oestrogen receptor, progesterone receptor and HER2 status reported by TCGA. All analyses were restricted to female, primary breast cancer (N=1078). Lymph node involvement status was also obtained for each patient (N positive = 463, N negative = 450).

### Akt gene signature

We built a gene signature for Akt activation based on genes identified by Creighton et al (ref). Principal component analysis was conducted on the expression of these genes to identify the pattern of gene expression which explained the greatest variance in the data. The first principal component was used as the Akt gene signature which was then split into high and low groups at the median for visualisation. The correlation between ISG15 expression and the Akt signature was assessed using linear regression for each breast cancer subtype with and without lymph node involvement separately.

### Akt activation from RPPA

pAkt measured by RPPA was available for a subset of the patients included above (N=322, N node positive = 156, N node negative = 166). The relationship between ISG15 and UBE2L6 expression and Akt activation was examined using linear regression for each subtype and in node positive and node negative cancer separately. The Benjamini-Hochberg adjustment was used to interpret the results while accounting for the effect of multiple testing. All analyses were carried out in R (version 3.6.0).

## Supplementary Figure Legends

**Fig.S.1**. (A) Kaplan-Meier plot of disease free survival association with mRNA levels of UBE2L6, separated into two groups with high (blue) or low (red) levels, in lymph node positive patients (n=744). (B) UBE2L6 mRNA levels correlation with disease free survival in lymph node negative patients (n=350). (C) Kaplan-Meier plot of cohort displayed in (A) classified by breast cancer subtype: luminal A (n= 823), luminal B (n=1013), HER2 (n=286) and basal (n=424). p-values are displayed in the top right corner of each plot.

**Fig.S.2**. (A) Representative images of colonies obtained in soft agar experiment at 20x. (B) Relative invasion. Equal amounts of cells were seeded, invasion ability was measured using 3-D matrigel scratch wound assay, photos were taken every 3h for up to 48h and invasion was determined as percentage of wound closed per time point. Bar graph shows the average invasion of the different clones versus control cells, at 12, 24 and 48h ±S.E.M; n=3. (C) Representative images of invasion assay at 12h, 24h and 48h.

**Fig.S3**. (A) Western blot of WT, crISG15 and crUBC8 cells non-transfected or transfected with a FLAG-ISG15 expression vector for 48h, and treated with EGF 10ng/ml for 10min. Activation of Akt was detected by immunoblotting the levels of pAktser473. (B) Western blot of WT and crISG15 cells treated with insulin, 0, 1, 10 or 100nM for 30min. Activation of Akt was detected by immunoblotting the levels of pAktser473. (C) Representative time-course analysis of Akt and ERK activation among the CRISPR clones treated with EGF 10ng/ml for 0, 2, 5, 10, 30 or 60min. Activation of Akt was detected by immunoblotting the levels of pAkt; activation of ERK by immunoblotting the levels of ppERK. (D) Quantification of the time course for pAktser473. Bars display the average value ±S.E.M of an n=4; p-value < 0.05 (*), p-value < 0.01 (**). (E) Western blot of the different clones blotted to analyse PTEN levels.

**Fig.S4**. Schematic illustration of the workflow for mass spectrometry-based analysis of ISGylome (left) and whole proteome expression levels (right), and the comparison between the specific hits obtained. Indicated clones were lysed in presence of 1% of SDS. Lysates used to determine the ISGylome were diluted 10-fold and subjected to an ISG15 immunoprecipitation using anti-ISG15 antibody bound to protein-G agarose beads. Pulldowns were digested with Trypsin in 2M urea, peptides were subsequently purified and analysed using a QExactive™ Plus Hybrid Quadrupole-Orbitrap™ Mass Spectrometer, protein identification and quantification was performed using MaxQuant software. Equal amounts of lysates used for determination of total protein expression were loaded into filters, and digested sequentially with Lys-C and trypsin. Peptides obtained from both digestions were purified and analysed using a QExactive™ Mass Spectrometer, protein identification and quantification was performed using MaxQuant software.

**Fig.S5. ISGylation of GDI2 reduces its interaction with Rabs**. (A) Bar graph shows label free quantification for GDI2 levels in whole proteome MS analysis. Bars represent the average LFQ values ±SD. (B) WB of the different clones blotted to analyse GDI2 levels. (C) Bar graph shows the LFQ values of GDI2, ISG15, Rab5A and Rab11 obtained in MS analysis of GDI2 pull-downs in the indicated clones transfected with MYC-DDK-GDI2. Bars graphs display the average value ±SD. ISG15, Rab5A and Rab11 enrichment was determined by normalization of their LFQ values by GDI2. (D) Bar graph shows the intensity of the modified peptide display in (F). ISGylated peptide enrichment was determined by normalization of the ISGylated peptide intensity divided by total GDI2 intensity. (E) Quantification of the pAkt levels in WT and crGDI2 cells treated with EGF for 10min. Bars show average values ±S.E.M; n=4, p-value < 0.01 (**). (F) Fragmentation spectra of GDI2 K-GG peptide. Cos-1 cells were non-transfected, transfected with MYC-DDK-GDI2, treated with vehicle for 48h or transfected with MYC-DDK-GDI2 and treated with IFN1b 250Pm for 48h and then subjected to a FLAG immunoprecipitation, trypsin digestion and peptide purification. Peptides were analysed with a Orbitrap Fusion™ Lumos™ Tribrid™ Mass Spectrometer. Identification of proteins and peptide modification was performed using MaxQuant software.

**Fig.S6. GDI2 ISGylation regulates EGFR translocation to the Golgi apparatus**. (A) Representative confocal immunofluorescence images, at 60x, of WT cells, crGDI2, crGDI2 transfected with GDI2wt and crGDI2 cells transfected with GDI2-KRtrip and stimulated with EGF 10ng/ml for 10min. The channels show EGFR (green), the Golgi marker GM130 (red), early endosome marker EEA1 (magenta), and nuclear marker, DAPI (blue). 10µm scale bars are displayed in the bottom-left corner. (B) Detailed view of EGFR and GM130 co-localization, visualisation of the co-localization between EGFR and the Golgi marker GM130 of the images displayed in (A) was performed by determination of co-localization areas using the Imaris software and a colocalization channel was built with it (yellow), DAPI (blue) is shown as a reference for the nucleus.

**Table S.1. Differentially expressed proteins in the different ISGylation clones**. Datasheet shows the FASP data of hits with differential expression versus WT cells. Differences in expression were determined and ratio versus the WT and as statistical changes using LFQ values normalized by protease values.

**Table S.2 Endogenous ISG15 pulldown MS analysis**. Datasheet 1 shows the specific hits for ISGylation. Determination of ISGylation was performed by enrichment of hits in samples from WT and crUSP18 cells versus the negative controls, crISG15 and crUBC8. Datasheet 2 shows the Maxquant obtained data without further analysis to show ISG15 levels. Datasheet 3 shows KEGG pathways clusters using the STRING database (www.string-db.org/). p-values, FDR, number of hits and list of hits are displayed in the table.

**Table S.3 Flag-GDI2 pulldown MS analysis**. Datasheet shows the specific hits for GDI2 interaction. Data was filter versus a negative control using the average LFQ, the obtained hits were normalized in each sample by their GDI2 LFQ values, and enrichment was performed to identify changes in interactome dependent of ISGylation status.

